# Evaluation of nafamostat mesylate safety and inhibition of SARS-CoV-2 replication using a 3-dimensional human airway epithelia model

**DOI:** 10.1101/2020.09.16.300483

**Authors:** D. Lynn Kirkpatrick, Jeffrey Millard

**Affiliations:** Ensysce Biosciences Inc. and Covistat Inc. La Jolla, CA 92037, www.ensysce.com

## Abstract

In the current COVID-19 pandemic context, Ensysce and its subsidiary Covistat have been working to repurpose nafamostat mesylate as an effective oral and inhalation treatment against SARS-CoV-2 infection. Prior reports used cell lines to demonstrate the antiviral potential of nafamostat against coronaviral infections and determined its mechanism of action through inhibition of transmembrane protease serine 2 (TMPRSS2). We selected a biologically relevant pre-clinical experimental model of SARS-CoV-2 lung infection using a 3D human reconstituted airway epithelial model of nasal origin to characterize the effects of nafamostat on tissue-level cellular ultrastructure and viral infection kinetics. Our results confirm the not only the relevance of this model for the preclinical evaluation of safety and efficacy of antiviral candidates, but also the highly potent nature of nafamostat SARS-CoV-2 antiviral activity. The studies described herein provided evidence demonstrating the therapeutic potential of nafamostat against COVID-19, as well as its safety upon exposure to lung airway cellular.

**One Sentence Summary:** Using a human airway model, study demonstrates the powerful inhibitory effect of nafamostat on SARS-CoV-2 genome copy detection when applied apically.

## INTRODUCTION

Nafamostat mesylate (herein referred to as nafamostat) is a synthetic, broad-specificity protease inhibitor of the classical and alternate pathways of complement activation. Nafamostat inhibits serine proteases such as F.VIIa, F.XIIa, kallikrein, thrombin, components of the complement system and trypsin. Nafamostat prevents coagulation abnormalities, and organ failure induced by complement-activated leukocytes in animal models of disseminated intravascular coagulation (DIC) (Cho et a. 2011; Hitomi and Fujji, 1982; Iwama et al. 2003). Nafamostat (Futhan^®^, Torii Pharmaceuticals) is an approved pharmaceutical for intravenous, intraarterial or extracorporeal use in Japan and Korea and has been used clinically since the 1980s (Honda et al. 2003). Nafamostat has not been approved for oral, inhalational or intranasal use in any jurisdiction.

Coronaviruses (CoV) are a group of highly diverse, enveloped, positive-sense, single-stranded RNA viruses that cause respiratory, enteric, hepatic and neurological diseases of varying severity in a broad range of animal species, including humans. Over the past 12 years, at least 3 novel βCoVs have emerged; severe acute respiratory syndrome CoV (SARS-CoV), Middle East respiratory syndrome CoV (MERS-CoV), and most recently SARS-CoV-2, responsible for coronavirus disease 2019 (COVID-19), which has caused an ever increasing number of cases and hospitalizations throughout the world. Nafamostat has been shown to inhibit viral entry in MERS and COVID-19 CoVs (Yamamoto et al., 2016; Hoffmann et al., 2020) and clinical trials with infusional nafamostat have been initiated in Asia.

Nafamostat’s inhibition of the proteolytic enzyme transmembrane protease serine 2 (TMPRSS2) which activates the S protein, causing infection, allows it to target the host and not the virus itself, explaining its broad utility against coronaviral infections. Ensysce had been exploring oral nafamostat for its ability to potently inhibit trypsin, in the development of overdose protection for trypsin activated abuse protected (TAAP) opioid prodrugs. These studies led to the current focus on the use of oral nafamostat to target the SARS-CoV-2 in the gut that results in many of the gastro intestinal symptoms of COVID-19, as well as intranasal nafamostat to prevent CoV viral entry in the human lung epithelia.

The safety and the efficacy of nafamostat were explored in the currently study, by its delivery to the apical surface of lung epithelia, using MucilAir™ technology (Epithelix, Sarl). The model is a pseudostratified and ready-to-use 3D model of human airway epithelium, constituted with primary human epithelial cells freshly isolated from nasal, tracheal or bronchial biopsies. When switched at the air-liquid interface, the progenitor cells undergo a progressive differentiation and polarization to a fully ciliated epithelia (**Figure 1**). The mature MucilAir™ is composed of basal cells, ciliated cells and mucus cells. The proportion of these various cell types is preserved compared to what one observes *in vivo* (Huang et al., 2011).

**Figure 1:**
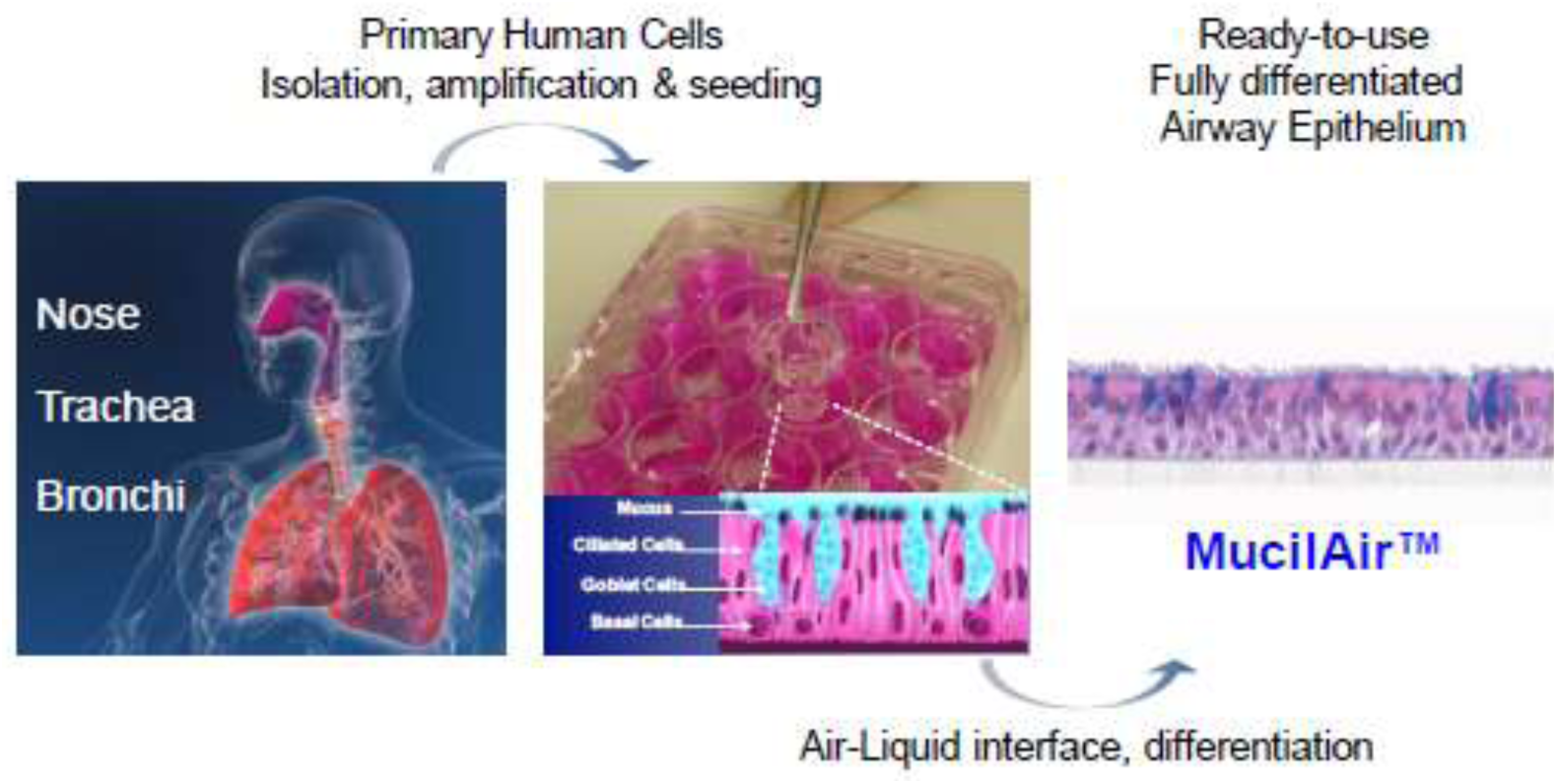
MucilAir™, a fully differentiated 3D in vitro cell model of the human airway epithelia. Epithelial cells were freshly isolated from the biopsies (nose), then seeded onto a semi-porous membrane (Costar Transwell, pore size 0.4 μm). After culture at air-liquid interface, the epithelia were fully differentiated, both morphologically and functionally.

This airway model is functionally differentiated, secretes mucus and is electrically tight (TEER>200 Ω.cm2). The activity of the main epithelial ionic channels, such as CFTR, EnaC, Na/K ATPase, is preserved and the epithelia is shown to respond in a regulated and vectorial manner to the pro-inflammatory stimulus, TNF-α (Huang et al., 2011). A large panel of cytokines, chemokines and metalloproteinases has been detected in the model (e.g. IL-8, IL-6, GM-CSF, MMP-9, GRO-α). Most importantly, it replicates the main function of the airway epithelial cells, the mucociliary clearance driven by synchronized cilia-beating.

The aim of this project was to evaluate antiviral effect of nafamostat upon SARS-CoV-2 viral infection (French circulating strain) using fully differentiated human nasal epithelial cells cultured at the air-liquid interface. Lung airway epithelia were reconstituted with a mixture of cells isolated from 14 different normal nasal donors, and the studies of the tissue integrity were carried out by Epithelix, Sarl.

The project was divided into two steps. Phase 1 involved a toxicological evaluation by examining tissue integrity, cytotoxicity, cilia beating and inflammatory reaction to repeated exposure to nafamostat. Phase 2 studied the effect of nafamostat on SARS-CoV-2 infection by measuring viral genome copy number after repeated dosing.

## MATERIALS AND METHODOLOGY

### Tissues (Epithelium source)

The lung airway model was comprised of a pool of human cells of nasal origin grown in batches as identified in Table 1. Table 2 provides the age and gender of each of the 14 samples that were used.

**Table 1:** MucilAir™-Pool batch information.

| Batch number | Age of the patient | Sex of the patient | Age of cultures | Special comments |
| --- | --- | --- | --- | --- |
| HF-MP0009<br>week 16 | - | - | 36 days<br>(ALI) | Pool of donors<br>No pathology<br>Phase 1 |
| HF-MP0009<br>week 21 | - | - | 35 days | Pool of donors<br>No pathology<br>Phase 2 |

**Table 2:** Nasal epithelia source for MP0009 batch

| Batch number | Age | Gender |
| --- | --- | --- |
| AB081301 | 52 | M |
| AB080901 | 24 | M |
| AB080801 | 36 | M |
| AB080601 | 32 | M |
| AB080301 | 58 | M |
| AB079801 | 44 | F |
| AB079701 | 55 | M |
| AB078001 | 55 | M |
| AB076201 | ND | ND |
| AB075502 | 54 | F |
| AB074501 | 53 | M |
| AB073801 | 32 | F |
| AB073501 | 57 | M |
| AB073101 | 58 | M |

### Test Article

**Nafamostat mesylate** (Katsura lot CS190701): 4-[(Aminoiminomethyl)amino]benzoic acid 6-(aminoiminomethyl)-2-naphthalenyl ester dimethanesulfonate was prepared in DMSO at concentrations of 2, 20, 200, 2000, 20000 nM and frozen until use.

### Phase 1

Phase 1 evaluated nafamostat effect on tissue integrity following three exposures at time 0, 24 and 48 hr. Each study was conducted in replicates of 3, **Figure 2**. Negative and positive controls were carried out in replicates of 6. Untreated or vehicle controls were used in replicates of 3 each. Positive controls included Triton X-100 (10 %, 50 µl apical, 24 hours; for cytotoxicity) N=3 and Cytomix (500 ng/ml TNFα, 0.2 mg/ml LPS, 1 % FCS; for inflammation) N=3. Endpoints were assessed at 24, 48 and 72 hr (days 1, 2, and 3).

**Figure 2:**
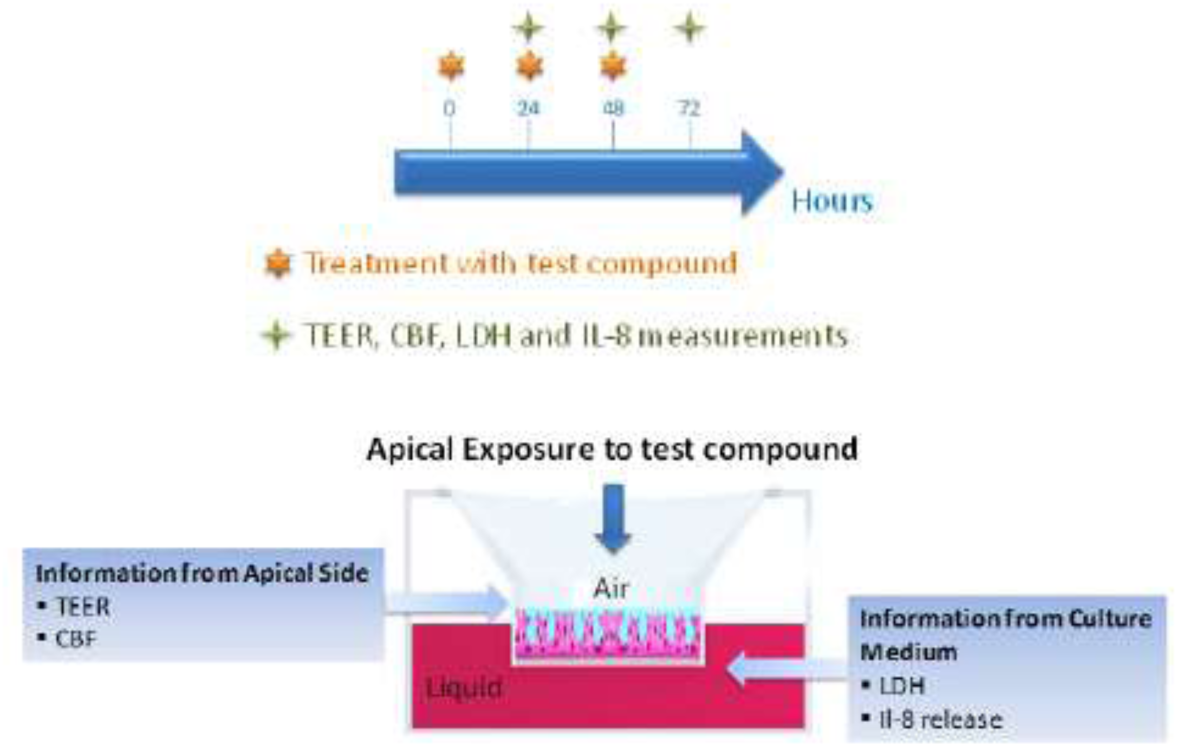
Apical exposure of test compound on epithelial airway model. End points were measured from apical side and basolateral culture medium.

### TEER

After addition of 200 µl of buffered saline/culture medium to the apical compartment of MucilAir™ cultures, resistance was measured with an EVOMX volt-ohm-meter (World Precision Instruments UK, Stevenage) for each condition. Resistance values (Ω) were converted to TEER (Ω.cm^2^) using the following formula: TEER (Ω.cm^2^) = (resistance value (Ω) - 100(Ω)) x 0.33 (cm^2^), where 100 Ω is the resistance of the membrane and 0.33 cm^2^ is the total surface of the epithelium.

### Cytotoxicity

Lactate dehydrogenase, a stable cytoplasmic enzyme that is rapidly released into the culture medium upon rupture of the plasma membrane was used to measure cytotoxicity caused by nafamostat. Basolateral medium (100 μl) was collected at each time-point and incubated with the reaction mixture of the Cytotoxicity Detection Kit^PLUS^ (LDH) following the manufacturer’s instructions (Sigma, Roche 04 744 926 001). The amount of the released LDH was quantified by measuring the absorbance of each sample at 490 nm with a microplate reader. To determine the percentage of cytotoxicity, the following equation was used (A = absorbance values):

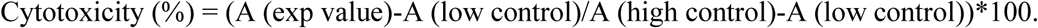

The positive control value was obtained by 10 % Triton X-100 apical treatment (24 hours). Triton X-100 causes a massive LDH release and corresponds to 100 % cytotoxicity. The negative controls (non-treated and vehicle) show a low daily basal LDH release, <5 %, which is due to a physiological cell turnover in MucilAir™.

### Cilia Beating Frequency (CBF)

Cilia beating frequency was measured by Epithelix using their in-house system consisting of three parts: a Mako G030B camera connected to a Zeiss Axiovert 200M microscope, a PCI card and a custom software. The cilia beating frequency expressed as Hz. 256 images were captured at high frequency rate (125 frames per second) at room temperature, cilia beating frequency was then calculated using validated Epithelix software.

CBF values may be subject to fluctuations due to parameters such as temperature, mucus viscosity or liquid applied on the apical surface of MucilAir™.

### Interleukin 8 ELISA assay

Pro-inflammatory cytokine Interleukin 8 (IL-8) was measured using ELISA assay (BD Biosciences 555244 (detection: 3-200 pg/ml) from basolateral medium of the airway cultures exposed to test compound. The assay was performed according to the manufacturer’s instructions. Lyophilized standard was reconstituted, aliquoted and stored at -80oC. Each ELISA plate contained a standard curve. Samples were diluted with the appropriate assay diluent at the appropriate dilution rate, washing steps were performed manually and absorbance was measured at 450 and 570 nm using Tecan Infinite 200.

### Phase 2

#### Virus Inoculation

The SARS-CoV-2 strain used in the study was isolated by directly inoculating VeroE6 cell monolayers with a nasal swab sample collected from Bichat Claude Bernard Hospital, Paris. Once characteristic cytopathic effect was observable in more than 50% of the cell monolayer, supernatants were collected and immediately stored at -80 °C. The complete viral genome sequence was obtained using Illumina MiSeq sequencing technology and was deposited under the name BetaCoV/France/IDF0571/2020. Viral stocks were titrated by tissue culture infectious dose 50% (TCID50/ml) in VeroE6 cells, using the Reed & Muench statistical method. These studies were conducted at VirNext (BSL3 facilities).

Prior to infection, the apical side of the airway cultures were washed twice for 10 min. Inoculations were performed with 150 μl at a theoretical multiplicity of infection (MOI) of 0.1 (50 000 TCID50 for an average of 500 000 cells in MucilAir™), applied to the apical side of the cultures for 1 hour at 37°C, 5% CO_2_. Non-infected control, Mock, was exposed also to 150 μl of culture medium on the apical side for 1 hour. Unbound viruses were removed after one hour of incubation period. New viral particles were collected by 10 min apical washes at 48 and 72 hours post-inoculation and quantified by RT-qPCR.

Reference antiviral drug remdesivir was purchased from MedChemExpress, (HY-104077) and was diluted in DMSO and used at 5 μM (final concentration of DMSO was 0.05 %) in the basolateral medium of separate cells. The reference antiviral was added after one hour of viral inoculation and changed every day.

#### Real-time Taqman probe RT-PCR

From the 200 μl apical washes, 140 μl was used for viral RNA extraction with the QIAamp® Viral RNA kit (Qiagen), obtaining 60 μl of eluted RNA. Viral RNA was quantified by quantitative RT-PCR (EXPRESS One-Step Superscript™ qRT-PCR Kit, Invitrogen, 1178101K) using 2 μl of viral RNA with Mastermix and two **ORF1b-nsp14** specific primers (5’-TGGGGYTTTACRGGTAACCT-3’; 5’-AACRCGCTTAACAAAGCACTC-3’) and probe (5’-FAM-TAGTTGTGATGCWATCATGACTAG-TAMRA-3’) **of SARS-CoV-2** designed by the School of Public Health/University of Hong Kong (Leo Poon, Daniel Chu and Malik Peiris). Samples were run on StepOnePlus™ Real-Time PCR System (Applied Biosystems). Ct data were determined and relative changes in gene expression were calculated using the 2^-ΔCt^ method and reported as the fold reduction relative to the mean of vehicle treated infected inserts.

#### Statistical analysis

Data were expressed as mean ± standard error of mean. Differences between three or more groups were tested by one-way ANOVA with Dunnett’s multiple comparison post-tests or nonparametric Kruskal-Wallis test with Dunn’s post-tests using Prism 6 GraphPad software (La Jolla, USA). Differences between two groups were tested by Student’s t test or nonparametric Mann-Whitney test. The values P<0.05 were considered statistically significant.

## RESULTS

### Phase 1: Evaluation of cytotoxicity of nafamostat on 3D airway epithelial model

#### Effect of Nafamostat on tissue integrity after repeated dosing

Nafamostat up to 20 µM did not cause any impairment of barrier function (tissue integrity limit was 100 ohm.cm^2^). The decrease of TEER at 2 and 20 µM suggested nafamostat caused an increase of ion channel activity (**Figure 3a**). TEER is a dynamic parameter that reflects the state of epithelia and is typically between 200 to 600 Ω.cm^2^. An increase of the TEER value reflects a blockage of the ion channel activities. A notable decrease of the TEER values (but > 100 Ω.cm^2^) could be observed in certain cases, reflecting an activation of the ion channels. Disruption of cellular junction or holes in the epithelia result in TEER values below 100 Ω.cm^2^. When an epithelium is damaged, a decrease of TEER would be associated with an increase of LDH release or a decrease of the cell viability.

**Figure 3:**
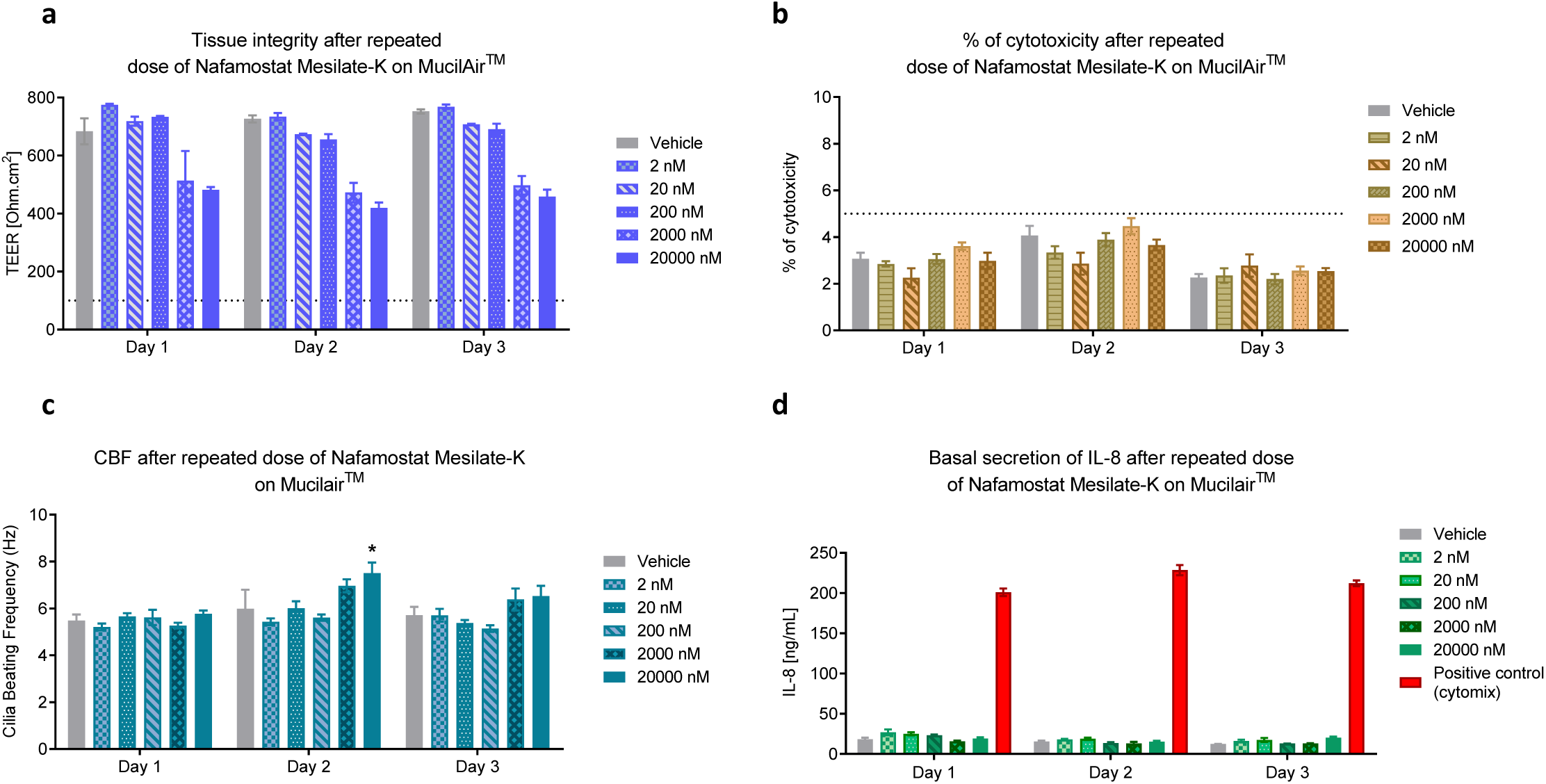
Effect of repeated (3 days) apical exposure to Nafamostat on MucilAirTM-Pool (n=3 cultures) measuring four outcomes. a) Transepithelial electrical resistance (TEER). Threshold limit value is 100 Ω.cm^2^. b) LDH release indicating cytotoxicity. Threshold limit value is 5 % cytotoxicity, which corresponds to a daily physiological LDH release in MucilAir™. c) Cilia beating frequency. d) Basal interleukin 8 secretion. Statistical comparison was performed using two-way ANOVA with Dunnett’s multiple comparison post-tests comparing all conditions to vehicle (Prism 6.0 GraphPad, *p<0.05, **p<0.01***p<0.001, ****p<0.0001).

#### Cytotoxicity of nafamostat after repeated dosing

There was no cytotoxicity observed following 3 days of dosing up to 20 µM, **Figure 3b**. Positive control 10% Triton X-100 solution induced toxicity was 100%.

#### Effect of nafamostat on Cilia Beating

The vehicle treated cultures showed cilia beating frequency of 5.5-6 Hz at room temperature, which is in the normal range of MucilAir™ (4-8 Hz). Exposure to nafamostat did not globally modify cilia motion. A small increase of CBF was observed at 2 µM at Day 2 and Day 3, and CBF was found to be significantly higher (20 µM Day 2), **Figure 3c**.

#### Effect of nafamostat on Interleukin 8

The vehicle treated cultures showed Interleukin 8 concentration of 12-18 ng/ml in the basal culture medium, which is in the normal range of MucilAir™ (5-50 ng/ml). Exposure to nafamostat did not modify basal IL-8 secretion, **Figure 3d**. Cytomix induced a significant increase of basal IL-8 concentration (p<0.0001).

### Phase 2: *Antiviral assessment of Ensysce Biosciences’ compound upon SARS-CoV-2 infection*

#### Relative virus genome copy

Exposure to Nafamostat showed dose dependent inhibition of apical genome copies of SARS-CoV-2 on the 3-D airway model. Exposure to nafamostat at 20 µM concentration strongly inhibited SARS-CoV-2 replication at both time points, with p<0.05 at 48 hr and p<0.01 at 72 hr, **Figure 4**. The magnitude of inhibition was 3.4 log at 48 hours and 6.2 log at 72 hours compared to vehicle. Exposure to nafamostat at lower concentrations of 2 and 0.2 µM also showed an antiviral activity on apical SARS-CoV-2 genome copies with an inhibition of 2.6 log at 48 hours and 4.3 log at 72 hours (2 µM) and 1 log at both time points (0.2 µM), compared to vehicle.

**Figure 4:**
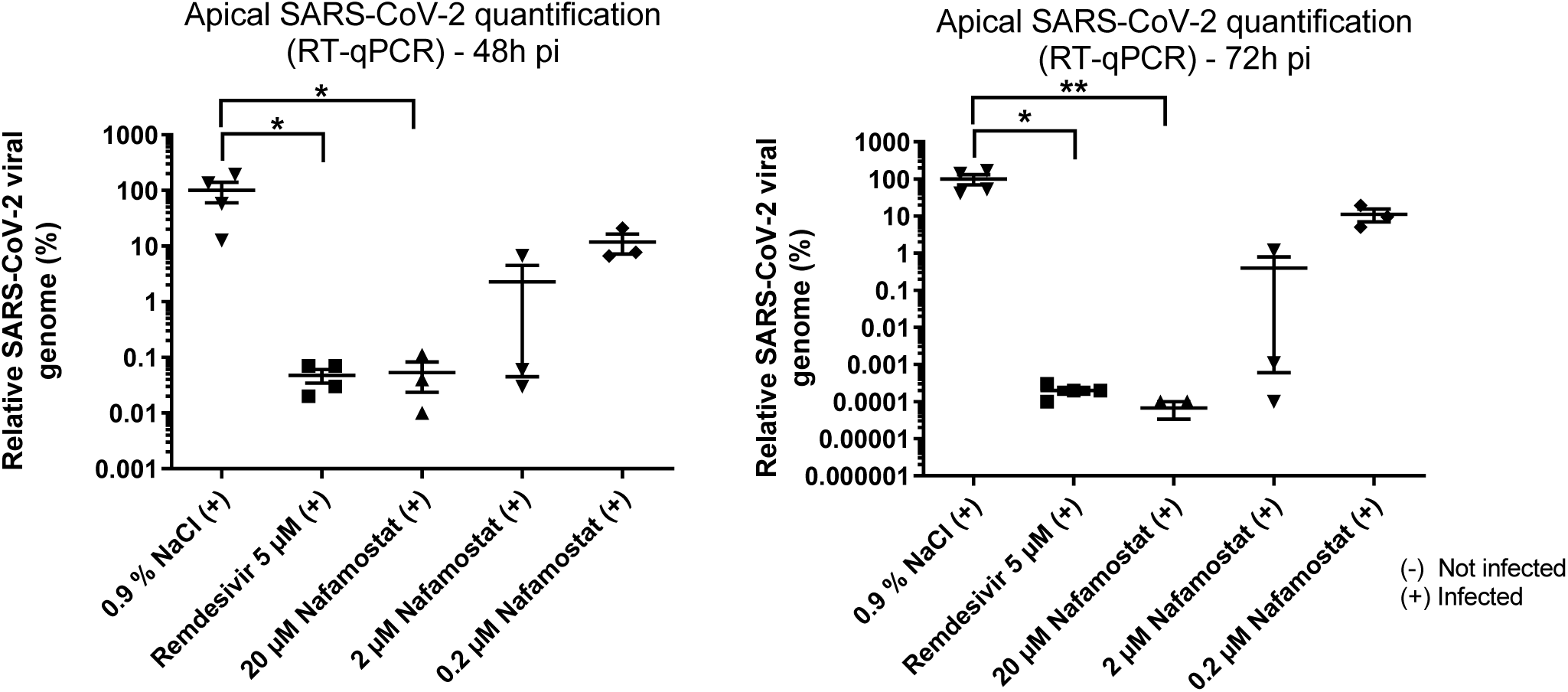
Relative SARS-CoV-2 genome copy number on the apical side of MucilAir™-Pool with one hour pre-treatment and repeated apical exposure of Nafamostat once a day (n=4, 3 cultures) at 48 hours (top) and 72 hours (bottom). Antiviral remdesivir at 5 *µ*M was added in basolateral medium at one hour post-inoculation. Statistical comparison was performed comparing all conditions to vehicle using Kruskal-Wallis test with Dunn’s posttests (Prism 6.0 GraphPad, *p<0.05, **p<0.01***p<0.001, ****p<0.0001).

Antiviral control remdesivir significantly reduced apical SARS-CoV-2 genome copies at both time points. The magnitude of inhibition was 3.4 log (mean Ct was 29) at 48 hours (vs. 0.9 % NaCl with a mean Ct of 18.3) and 5.7 log (mean Ct was 31.9) at 72 hours (vs. 0.9 % NaCl with a mean Ct of 13.2).

## DISCUSSION

The aim of this study was to evaluate nafamostat on Severe Acute Respiratory Syndrome Coronavirus 2 infection of *in vitro* respiratory epithelium. The epithelia used were of nasal origin from 14 donors, in a platform called MucilAir™-Pool. Initial evaluation of human airway epithelial cells potential toxicity by nafamostat demonstrated that there was no disruption of the tissue integrity at any of concentrations tested, by any of the methodology used. Modest ion channel activity was observed along with an increase in ciliary beating the two highest concentrations. These effects are considered to be beneficial for removal of mucoid material from lungs.

The study to evaluate the effects of nafamostat on SARS-CoV-2 infection, epithelia were pre-treated (1 hour) and retreated with nafamostat on the apical side once a day every day after respiratory virus infection. This treatment simulates the condition of using an inhalation nafamostat drug product for COVID-19 treatment. Quantification of effect of nafamostat on apical viral replication was measured by RT-PCR (genome copy number).

Although prior reports had suggested SARS-CoV-2 to cause a decreased TEER (Pizzorno et al., 2020), in this study SARS-CoV-2 did not modify tissue integrity, **Figure 5**.

**Figure 5:**
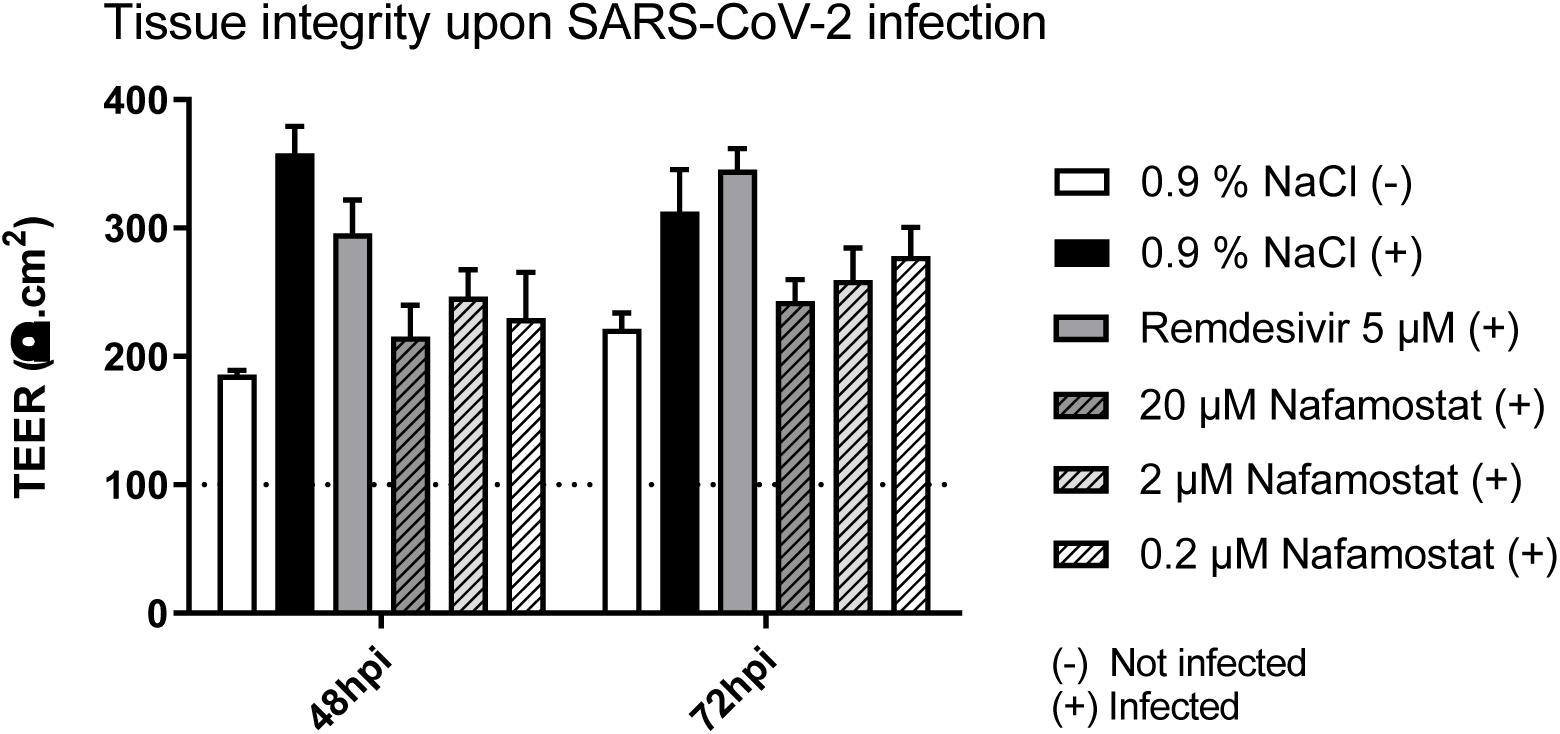
Tissue integrity as measure by TEER (Ω.cm^2^) with SARS-C0V-2 infection and treatment with nafamostat (0.2, 2.0 and 20 µM) or remdesivir (5 µM).

Exposure of the apical surface of the 3-D airway model to nafamostat shows dose dependent inhibition of apical genome copies of SARS-CoV-2. This data supports that reported by others who elucidated the mechanism of action of nafamostat on SARS-CoV-2 using cell lines. Nafamostat, an ultrapotent protease inhibitor, inhibits TMPRESS2, and here was shown to reduce viral entry over a range of concentrations. A decrease in viral copy of 1 log was observed at concentrations as low as 0.2 µM. A dose and time dependent effect was observed, with nafamostat (20 µM) at 72 hr providing a greater reduction of viral copy than that produced by remdesivir.

This study demonstrates both the safety and the powerful inhibitory effect of nafamostat when applied apically on human airway epithelia on SARS-CoV-2 genome copy detection.

